# CLOCK Mediates Delay in Cardiac Myofibroblast Formation and Attenuates Progression of Myocardial Fibrosis through Inhibition of Smad 3 Transcriptional Activity

**DOI:** 10.1101/2023.08.27.555032

**Authors:** Yongqiao Zhou, Minqi Liao, Lihua Chen, Yongzhao Yao, Zhichao Huang, Yanhua Yang, Shaohui Su, Guangzhu Liang, Zhihao Wu, Shumin Ouyang, Suxia Guo

## Abstract

**Introduction:** Myocardial fibrosis, characterized by excessive extracellular matrix deposition, leading to adverse cardiac remodeling and impaired function. The differentiation of cardiac fibroblasts into myofibroblasts plays a pivotal role in this process. The involvement of the core clock protein CLOCK in the formation of cardiac myofibroblasts and the advancement of myocardial fibrosis is not well understood. This study aims to investigate how CLOCK modulates cardiac myofibroblast differentiation and attenuates myocardial fibrosis by inhibiting Smad3 transcriptional activity.

**Methods:** In vivo, adeno-associated virus 9 was utilized for overexpression or silencing of the CLOCK gene via mice intravenous injection. In vitro experiments were conducted using primary cultures of cardiac fibroblasts isolated from mouse hearts to examine the expression levels of molecular markers associated with myofibroblast differentiation, such as α-smooth muscle actin (α-SMA) and collagen I. Additionally, a Smad3 luciferase reporter gene experiment was employed to assess the transcriptional activity of Smad3 under CLOCK regulation.

**Results:** The results demonstrate that CLOCK overexpression significantly reduces the expression of α-SMA and collagen I in cardiac fibroblasts, indicating a suppression of myofibroblast formation. Conversely, CLOCK knockdown enhances the expression of these fibrotic markers. Interestingly, CLOCK was found to negatively regulate the transcriptional activity of Smad3, suggesting a potential mechanism by which CLOCK exerts its inhibitory effect on cardiac fibrosis. Specifically, CLOCK interacts with Smad3, inhibiting its nuclear translocation and subsequent activation of fibro genic gene expression.

**Conclusion:** These findings highlight the crucial role of CLOCK in the regulation of cardiac myofibroblast formation and the progression of myocardial fibrosis. CLOCK appears to exert an inhibitory effect on fibrotic signaling pathways, partially through the inhibition of Smad3 transcriptional activity. This study provides novel insights into the molecular mechanisms underlying cardiac fibrosis and emphasizes the therapeutic potential of targeting CLOCK-mediated pathways for treating myocardial fibrosis.

## Introduction

Post-myocardial infarction (MI) cardiac fibrosis plays a critical role in the persistent and progressive remodeling of the heart^1–5^. Post-MI cardiac fibrosis is often accompanied by cardiac systolic and diastolic dysfunction, arrhythmias and adverse cardiovascular events^6,7^. However, the molecular mechanisms underlying this process are not fully elucidated. A comprehensive understanding of the molecular mechanisms involved in cardiac fibrosis is essential for the development of novel therapeutic strategies. The current treatment approach is still in the experimental phase, requiring further investigation and clinical trials to establish its efficacy and safety. Recent studies have uncovered the multifaceted and intricately regulated nature of cardiac fibrosis, predominantly driven by the transdifferentiation of fibroblast cardiomyocytes into myofibroblasts^8–14^.

Fibroblast cardiomyocytes play a crucial role in maintaining the balance of extracellular matrix (ECM) components. The ECM mainly consists of fibrous collagen, with type I collagen crucial rigidity and tensile strength, preserving the structural integrity of the ventricular wall. Conversely, type collagen constitutes approximately 10% of the total myocardial collagen, contributing to the elasticity of the cardiac extracellular matrix. These collagens serve as architectural frameworks for myocardial tissue and are involved in continuous synthesis and degradation processes^15–18^. Maintaining the appropriate distribution of collagen within the myocardial interstitium is vital for the preservation of cardiac function and myocardial tissue integrity. Cardiac fibrosis occurs when the delicate balance between collagen synthesis and degradation shifts toward increased collagen deposition and expansion of the ECM^19–23^.

Activation of the Smad 3 pathway plays a crucial role in the development of cardiac fibrosis, representing a significant pathological change^24–30^. Through the initiation of Smad 3 signaling, cardiac fibroblast differentiation can drive the progression of cardiac fibrosis^31,32^. Moreover, Smad 3 involved in the regulation of extracellular matrix protein synthesis, integrin transcription, and upregulation of α-SMA^33–35^. Notably, the activation of Smad 3 within the infarcted area of the heart is essential for the formation of tightly-knit myofibroblasts^36–38^, which are vital for maintaining the structural integrity of the infarcted ventricle. As a key component of the circadian machinery^39–41^, the CLOCK protein plays a vital role in regulating various aspects of cardiometabolic function, contractility, and the rhythmicity of genes and proteins^42–44^. Although the circadian rhythm has a significant impact on the timing of adverse cardiovascular events^42–45^, there are still unanswered questions regarding the detrimental effects of CLOCK disruption on cardiac structure and function, as well as its contribution to the development of age-related heart disease.

The regulatory effects mentioned above have brought CLOCK into the forefront of clinical investigation. However, there have been no reports to date that explore the involvement of CLOCK in cardiac fibrosis or its potential regulation by Smad 3. Therefore, the objective of this study was to examine the role of CLOCK and elucidate the underlying molecular mechanisms in cardiac fibrosis. To achieve this, we utilized myocardial tissue from a mouse model of post-myocardial infarction fibrosis and observed reduced levels of CLOCK both cardiac fibroblasts and fibrotic regions. Subsequent experiments were conducted to investigate the impact of CLOCK on inhibiting the differentiation of cardiac fibroblasts into myofibroblasts in vivo and cardiac fibrosis in vitro. Our aim was to validate CLOCK’s protective effect against myocardial infarction and explore its interaction with Smad 3. Overall, our findings reveal the crucial involvement of CLOCK in myocardial infarction and highlight its potential as a novel therapeutic target for patients with post-MI myocardial fibrosis.

## Methods

### Animal Model Establishment and Tissue Collection

Male C57BL/6J wild-type mice, aged 8 weeks. were procured from Beijing Weitong Lihua Co., Ltd. and were carefully chosen for this study. To comply with the ethical requirements for handling experimental animals set by Dongguan People’s Hospital, the mice were housed in a controlled environment with an SPF-grade feeding area at Beijing Weitong Lihua Co., Ltd. The mice were subjected to a 12-hour light-dark cycle and provided with ample water and feed.

To achieve overexpression of CLOCK in the mice’s heart, we utilized serotype 9 adeno-associated virus (AAV9). A specific number of 8-week-old male C57BL/6J mice were randomly selected according to the experimental design, and the AAV virus carrying the CLOCK gene was injected into their tail veins. This injection was performed for a duration of 4 weeks to ensure stable infection by the AAV virus.

Once the mice were adequately prepared for experimentation, they were anesthetized using isoflurane. Blood samples were collected from the orbital venous plexus and left at room temperature for 20 minutes. After centrifugation at 3000 rpm and 4°C, serum was isolated and stored at −80°C for further analysis. The chest cavity was promptly opened, and the heart was perfused with pre-cooled phosphate-buffered saline (PBS). Subsequently, the heart was removed, weighed while still moist, and photographed. Certain portions of the heart were fixed in a 4% paraformaldehyde solution for the creation of paraffin sections. Another part of the heart was preserved in a frozen storage tube, rapidly submerged in liquid nitrogen, and then transferred to −80°C to ensure long-term storage.

### Echocardiographic Assessment

The echocardiographic evaluation of the thoracic cardiac region in mice was performed four weeks after myocardial infarction (AMI) using the state-of-the-art Vevo2100 high-resolution imaging system. To ensure optimal imaging conditions, specialized depilating cream was used to gently remove mouse hair. The mice were efficiently anesthetized with isoflurane, and then the ultrasound probe was accurately positioned on the left anterior chest wall to obtain detailed two-dimensional images depicting the long axis of the heart. Three consecutive cardiac cycles were recorded and thoroughly analyzed for each experimental group. A meticulous focus was placed on acquiring M-mode echocardiographic images, which were indispensable for precise quantification of various cardiac parameters, including the left ventricular wall thickness (LVAWs, LVAWd, LVPWs, LVPWd), the left ventricular systolic and diastolic internal diameters (LVSDd, LVEDd), left ventricular ejection fraction (LVEF%), left ventricular short axis shortening fraction (LV fractional shortening%), left ventricular cardiac output (LV cardiac output), and left ventricular output per stroke (LV stroke volume).

### HE Staining

The HE staining procedure was meticulously conducted following the recommended methodology outlined in the HE kit. Three sections of each paraffin-embedded specimen underwent a careful and detailed series of steps to achieve optimal transparency. The nuclei were gently treated with hematoxylin for duration of five minutes, followed by a thorough rinsing in distilled water for ten minutes. Subsequently, a fresh batch of distilled water was briefly used for a ten-second rinse. In order to enhance the intensity of the stain, a precise application of 95% ethanol was conducted for a mere five seconds. Then, with meticulous precision, the eosin staining solution was skillfully applied for just thirty seconds, immediately followed by a two-minute immersion in 95% ethanol. An additional two-minute immersion in a fresh batch of 95% ethanol was performed. The achievement of transparency was diligently accomplished by carefully exposing the specimens to xylene for a period of five minutes, promptly followed by a repetition of this process using clean xylene for another five minutes. Finally, the meticulously treated specimens were protected with a layer of neutral gum. This resulted in the nuclei assuming a vibrant blue hue, while the cytoplasm exhibited a delicate pink shade.

### Masson Staining

Masson staining was performed following the recommended protocol provided by Solarbio Company. Three sections from each specimen underwent to standard dewaxing and dehydration procedures. The cells were stained using a compatible Weigert ferricatoxylin staining solution for a duration of 5 minutes. Subsequently, the acid ethanol differentiation solution was applied for 10 seconds, followed by double steaming washes. Masson’s staining solution was then used to stain the samples for 3 minutes, followed by rinsing with double-steamed water. An additional double steaming step was performed for 1 minute. To achieve Lichunhong staining, the samples were exposed to the staining solution for 5 minutes. A two-minute wash with phosphomolybdic acid followed, followed by a one-minute wash with a weak-acid working solution. Aniline blue staining solution was then applied for 1 minute, followed by another one-minute wash with the weak-acid working solution. Rapid dehydration was carried out using 95% ethanol, with three consecutive ten-second immersions in anhydrous ethanol. Subsequently, xylene was used three times for 1 minute each to achieve transparency. Finally, neutral gum sealing sheets were applied to fix the samples, and they were observed under a microscope. As a result of the staining, collagen tissue appeared blue, while the muscle fibers and red blood cells were stained red.

### Sirius Red Staining

For the Sirius red staining, we followed the recommended protocol providedby Solarbio Company. Three sections were carefully chosen from each specimen and undervent standard dewaxing and dehydration procedures. Subsequently, the slides were treated the Weigert Iron-hematoxylin staining solution for a duration of 10 minutes. Afterward, the slides were rinsed with tap water for 5 minutes and subjected to double steaming for 10 seconds to ensure proper staining conditions. The Sirius red staining solution was then applied and allowed to incubate for 1 hour. Following this, incubation period, the slides were thoroughly washed with distilled water, repeating the process three times to remove any excess stain. After routine dehydration, the sections were made transparent and mounted with neutral gum sealing sheets. Finally, the samples were observed under a microscope. Remarkably, the collagen fibers exhibited a vibrant red coloration, highlighting their presence and distribution within the tissue. Nuclei appeared blue, providing crucial information about cellular morphology, and muscle fibers displayed a yellow hue, aiding in the identification and characterization of muscular components.

### Cell Culture and Treatment

For this study, C57BL/6J mice aged between 1 and 3 years were included. Thorough disinfection of the thoracic and abdominal regions was performed using a 75% ethanol solution. The hearts were carefully extracted, ensuring complete removal of all connective tissue and adipose deposits. Subsequently, the mouse ventricles were finely minced into tissue blocks using a tissue digestive fluid consisting of 0.25% trypsin and 0.25% collagenase. The digestion process was carried out in a temperature-controlled water bath shaker set at 37℃ with an oscillation rate of 80 revolutions per minute. After 20 minutes, the centrifuge tube was removed, and the digestion was stopped by adding DMEM supplemented with 10% FBS and 1% double-antibody. The resulting cell suspension underwent centrifugation and was then resuspended in a complete growth medium. The cell resuspensions were seeded onto plates and placed in a controlled incubation environment at 37℃ and 5% CO₂ for 90 minutes. Following this initial incubation, the medium was replaced with fresh complete medium once at the 48-hour mark. The primary cardiac fibroblasts (CFs) obtained from these cultures were subsequently passaged at a ratio of 1:3. For experimental investigations, cells from the third generation were utilized.

### Cell Synchronization

In order to synchronize cultured myocardial fibroblasts, lactating rats were utilized as the source for these cells. The myocardial fibroblasts extracted and placed in a nutritive-rich solution supplemented with 10% fetal bovine serum (FBS). The cells were then carefully distributed into the appropriate number of dishes, ensuring an equal number of cells in each well. When the cell confluence reached approximately 80%, the medium was replaced with a specific formulation containing 50% equine serum. The purpose of this serum change was to synchronize cellular activity for a duration of two hours. Following this synchronization phase, the medium was subsequently replaced with DMEM medium containing 0.4% FBS, which facilitated the subsequent experiments. It is important to note that the initiation of the high-concentration equine serum treatment was designed as the zero time point for cell synchronization.

### Western blot analysis

To assess protein expression, Western blot analysis was conducted following established protocols. The membranes were incubated overnight at 4°C with primary antibodies, including anti-SMA (ab9725, 1:1000, Abcam), anti Smad3 (ab76836, 1:1000, Abcam), anti-Collagen I (10879-1-AP, 1:1000, Proteintech), anti-Collagen III (10882-1-AP, 1:1000, Proteintech), and anti-GAPDH (sc-365062, 1:3000, Santa Cruz Biotechnology). Following primary antibody incubation, the membranes were further incubated at 37°C for 1 hour with horseradish peroxidase-conjugated secondary antibodies: goat anti-rabbit (sc-2004, 1:5000, Santa Cruz Biotechnology) or goat anti-mouse (sc-2005, 1:5000, Santa Cruz Biotechnology). Immunoreactive bands were visualized using Immobilon Western Chemiluminescent HRP Substratet (Amersham Biosciences U.K. Ltd), and band intensities were quantified using Image-Pro Plus 6.0 software. Densitometry analysis was performed on three independent experiments, and the relative expression level of the target proteins were normalized to the intensity of the GAPDH band.

### Quantitative Real-Time PCR (qPCR)

RNA extracion from the cells and mouse hearts was performed using the sophisticated Trizol reagent (Takara, Kyoto, Japan). Subsequently, the RNA was subjected to reverse transcription to generate RNA cDNA using the widely recognized PrimeScriptRT kit (Takara, Kyoto, Japan) provided by the esteemed Bio-Rad. Accurate quantification of mRNA levels was meticulously conducted by employing the GAPDH reference gene and utilizing the cutting-edge SYBR Green kit (Takara, Kyoto, Japan) on the state-of-the-art Real-time PCR Detection System manufactured by Bio-Rad. For specific details regarding the primer sequences used, please refer to Supplementary Table 1, which provides comprehensive information on the primers employed in this study.

### Transfection and lentiviral infection of myocardial fibroblasts and HEK-293T cells

For transfection, the Lipofectamine RNAiMAX reagent (Invitrogen) was used, following the manufacturer’s instructions and specific concentrations requirements. Special attention was given to the meticulous transfection of siRNA into myocardial fibroblasts, ensuring optimal delivery and efficacy. Plasmids were also prepared with careful attention to detail using Lipofectamine 3000 regent, following the manufacturer’s instructions, and subsequently transfected into cultured VSMC or HEK-293T cells. The effectiveness of cell transfection was diligently evaluated using real-time polymerase chain reaction (PCR) methodologies.

To achieve overexpression of the CLOCK gene in cardiac fibroblasts, cells were cultivated until they reached a density of 70-80%. Recombinant lentiviral particles carrying CLOCK shRNA (obtained from Santa Cruz Biotechnology, sc-108080) were then introduced into the cardiac fibroblasts. As a control, CLOCK-sh-Scr lentiviral particles were also added to the cells, ensuring the rigorous and meticulous experimental design.

### Isolation and Culture of Cardiac Fibroblasts

Cardiac fibroblasts were obtained and cultured from suckling C57BL/6J mice. To ensure sterility, the newborn mice were subjected to sterilization using a 75% ethanol solution within 1-3 days of birth. Subsequently, the murine hearts were carefully extracted, ensuring the removal of surrounding connective tissue and adipose matter. The ventricles of the hearts were meticulously sectioned into tissue blocks and immersed within a tissue digestion compound consisting of 0.25% trypsin and 0.25% collagenase. The tissue digestion process was carried out in a constant bath oscillator operating at a consistent speed of 80 revolutions per minute, and the temperature was maintained at 37℃. After 20 minutes of enzymatic breakdown, the centrifuge tubes were carefully retrieved, and the digestion process was promptly terminated by the adding DMEM supplemented with 10% fetal bovine serum and 1% dual antibody solution. The resulting cellular suspension was then subjected to centrifugation, and the pellet obtained was resuspended in a comprehensive medium. These cell suspensions were incubated within a controlled environment of an incubator set at a temperature of 37℃, enriched with 5% carbon dioxide gas, for a duration of 90 minutes. Subsequently, the medium was replaced with a fresh, complete medium. Following a 1:3 dilution of primary myocardial fibroblasts, the third generation of cells was carefully selected for subsequent experimental investigations.

### ChIP assay

Primary fibroblasts from milk rats were transfected with Adenovirus-mediated overexpression of CLOCK. To establish cross-linking, The cells were fixed using 0.2 ml of a 10% formaldehyde solution (Sigma, Ontario, Canada) to. Subsequently, the fixed cells were centrifuged at 1400 g for 5 minutes at 4°C. The resulting pellets were then reconstituted in 1X PBS supplemented with a protease inhibitor cocktail (PIC), and underwent another round of centrifugation under the same conditions. After centrifugation, the pellets were resuspended in a low-salt buffer and subjected to sonication.

For immunoprecipitation, anti-CLOCK antibodies or normal mouse serum (Cell Signaling, Massachusetts, USA) were allowed to incubate with protein G Dynabeads (Life Technologies, Ontario, Canada) for 6 hours. Then, DNA samples were added to the antibody/Dynabead mixtures, followed by overnight incubation at 4°C. Subsequently, the samples were washed with a salt buffer to remove non-specific binding. The resulting eluted samples were subjected to reverse cross-linking through an overnight incubation at 65°C. DNA fragments were purified using phenol: chloroform extraction method and subsequently analyzed using quantitative polymerase chain reaction (qPCR). Please refer to for the primer sequences utilized in the PCR analysis Supplementary Table 2.

### Statistical analysis

All data analyses in this chapter were performed using the SPSS 23.0 software (SPSS Inc.). Descriptive statistics for the measurement data are presented as mean values with accompanying standard deviations. The compare two groups with normally distributed and equally variant measurement data, a one-way ANOVA was conducted, followed by pairwise comparison using the Bonferroni method. Furthermore, comparisons between the multiple groups were also conducted using a one-way ANOVA with the Bofferoni method employed for pairwise comparisons. To assess non-normal distributions across multiple groups, the chi-square test was applied. Statistical significance was defined as P <0.05 (* or #). All statistical analyses and data visualization were conducted using GraphPad Prism 8.0.

## Results

### Reduction of CLOCK protein associated with myocardial fibrosis development

To investigate the association between CLOCK protein and myocardial fibrosis, mice underwent cardiac echocardiography examination four weeks after acute myocardial infarction (AMI). The results revealed a significant decrease in left ventricular ejection fraction (LVEF) and left ventricular fractional shortening (LVFS) in the MI group compared to the Sham group (*p*<0.01), as presented in Supplementary Figure 1A-B, confirming the successful establishment of the AMI model. Serum B-type natriuretic peptide (BNP) is a crucial clinical measure for assessing heart failure. Using enzyme-linked immunosorbent assay (ELISA), we quantified serum BNP levels in mice and found a significant increase in the MI group compared to the Sham group (Supplementary Figure 1A-B) (*p* <0.01). To investigate the expression of CLOCK protein in mouse cardiac tissue(Figure 1A), Western Blot analysis with corresponding grayscale statistics demonstrated a notable elevation in the protein expression levels of alpha-smooth muscle actin (α-SMA) and Collagen I in the MI group compared to the Sham group (Figure 1B-C).

**Figure 1:**
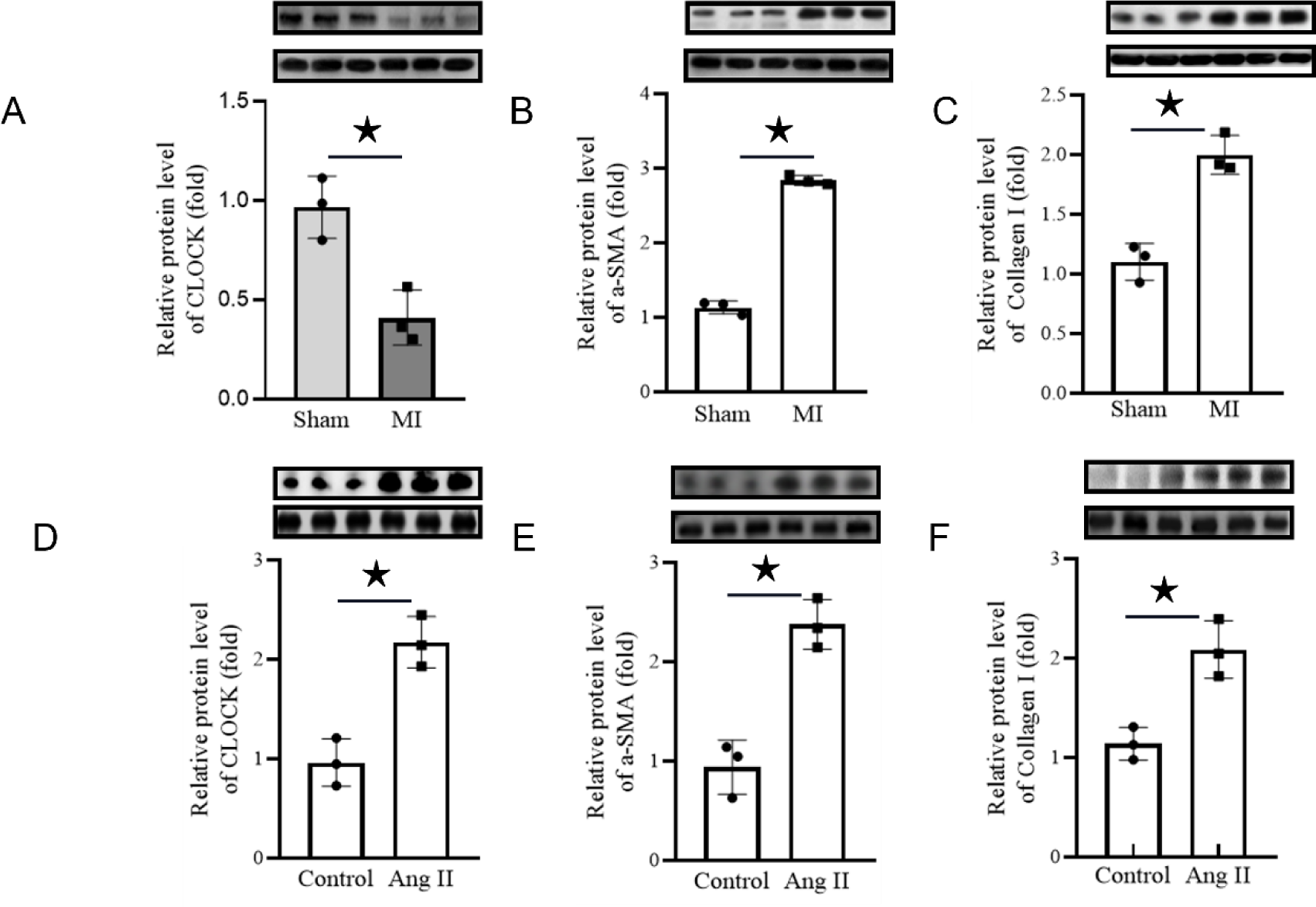
CLOCK Protein expression in Myocardial Fibrosis and Myofibroblasts. A-C:Four weeks post-acute myocardial infarction (AMI), the cardiac tissues were procured, and the protein expression profiles of CLOCK, alpha-smooth muscle actin (α-SMA), and Collagen I were assessed via western blot analysis.N = 3; *P<0.05. D-F:10-6M AngII stimulated mouse primary cardiac myocardial fibroblasts induced differentiation into myofibroblasts, and the protein expression levels of CLOCK, α-SMA, Collagen I were determined by western blot。N = 3; *P<0.05.

Previous studies have shown that exposing cardiac fibroblasts to a concentration of 10-6M angiotensin II (AngII) for 24 hours promotes their differentiation into myofibroblasts^46–50^. Interestinly, Western blot assays revealed a significant decrease in the expression of CLOCK protein within myocardial fibroblasts following AngII intervention (Figure 1D and Supplementary Figure 1C). In contrast, a substantial upregulation in α-SMA, Collagen I protein levels was observed compared to the control group (Figure 1E-F).

### CLOCK hinders the differentiation of mouse primary cardiac muscle fibroblasts into myofibroblasts in vitro

To investigate the impact of CLOCK knockdown on phenotypic differentiation, we examined suckling mouse primary cardiac fibroblasts. In vitro experiments demonstrated that si-RNA-mediated knockdown of CLOCK, followed by induction with 10-6M angiotensin II (AngII) for 24 hours, led to the differentiation of cardiac fibroblasts into myofibroblasts. Real-time PCR analysis confirmed a significant reduction in CLOCK mRNA expression in mouse primary myocardial fibroblasts upon si-CLOCK treatment (Figure 2A). Furthermore, protein immunoblot analysis revealed a decrease in CLOCK protein expression (Figure 2B), accompanied by an increase in levels of alpha-smooth muscle actin (α-SMA) and Collagen I proteins (Figure 2C-D), These findings suggest that CLOCK knockdown promotes the differentiation of primary myocardial fibroblasts into myofibroblasts.

**Figure 2:**
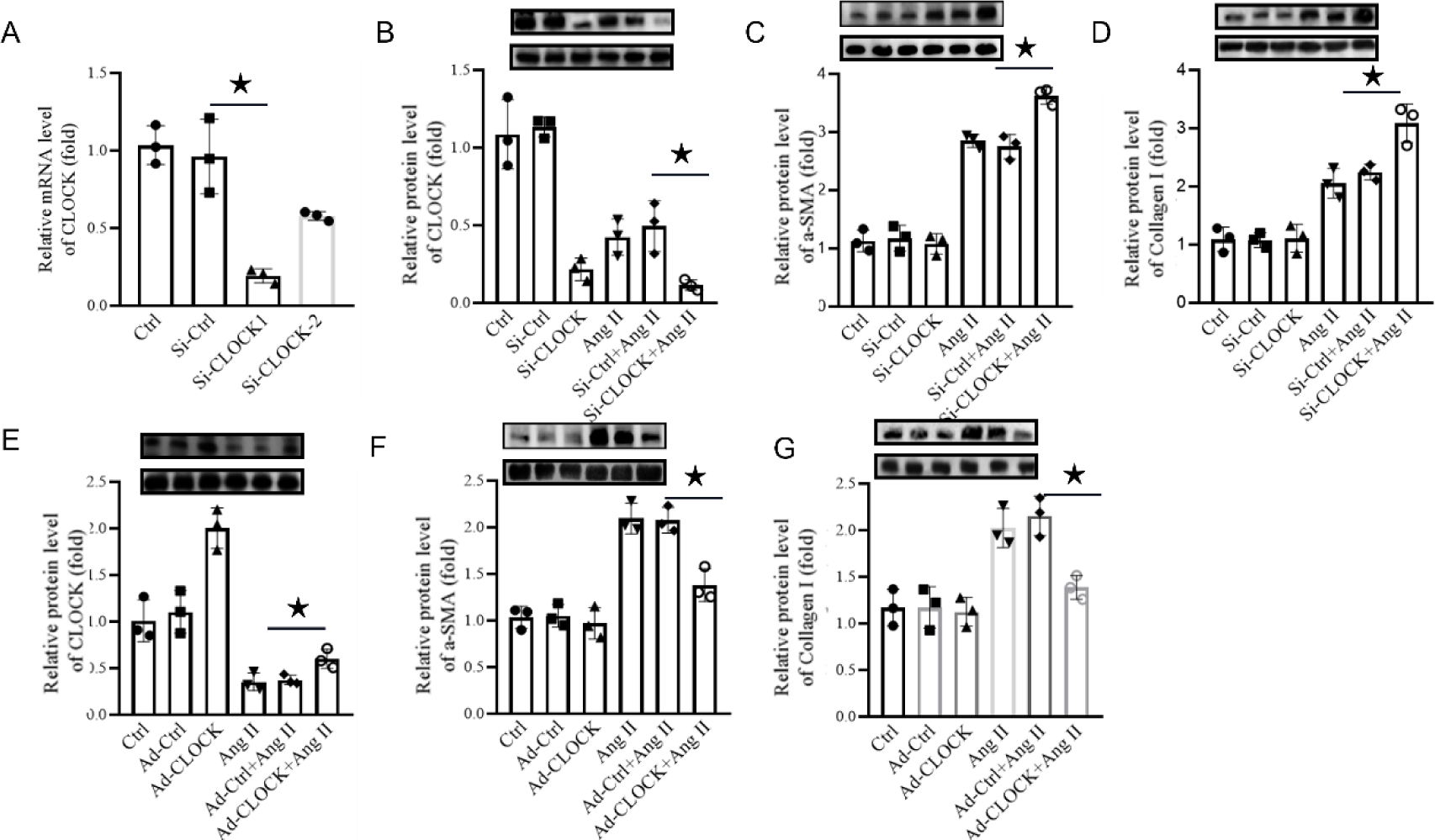
Overexpression of CLOCK in vitro inhibited the Ang-induced differentiation of cardiac muscle fibroblasts into myofibroblasts. A: CLOCK expression in Si-RNA knockdown cells, and CLOCK mRN levels in cardiac fibroblast cells were measured by Real-time PCR. B-D: Si-CLOCK knockdown cardiac fibroblasts under AngII induction; B-D: western blot for A-SMA and Collagen I.N = 3; *P<0.05。 E-G: the protein expression level of lentivirus mediated overexpression of CLOCK in cardiac fibroblasts and the induction of CLOCK, A-SMA, and Collagen I were measured by western blot.N = 3; *P<0.05, ***P<0.01。

Surprisingly, in vitro infection of primary cardiac fibroblasts with adenovirus Ad-CLOCK exhibited a striking phenomenon. These cells displayed an overexpression of the CLOCK gene while being exposed to Angiotensin II for 24 hours. Western blot analysis demonstrated a substantial down regulation of CLOCK expression (Figure 2E). Additionally, the levels of α-Smooth Muscle Actin (α-SMA), Collagen I proteins were found to be notably reduced (Figure 2F-G). These compelling results indicate that increasing CLOCK expression in cardiac fibroblasts inhibits the differentiation of Angiotensin-induced Cardiac Fibroblasts (CFs) into myofibroblasts, potentially mitigating the severity of cardiac fibrosis.

### CLOCK overexpression alleviates the severity of cardiac fibrosis in mice in vivo

To establish this, we employed serotype 9 adeno-associated virus (AAV9) to deliver a plasmid carrying the CLOCK gene, resulting in an animal model characterized by elevated CLOCK levels. Careful analysis and quantification of CLOCK protein in the cardiac tissue of mice were conducted. Remarkably, Real-time PCR and Western blot analyses, along with statistical analysis, demonstrated a significant increase in CLOCK expression in the AAV-CLOCK experimental group compared to the AAV-vehicle control (Figure 3A-B). Furthermore, the AAV-CLOCK cohort exhibited when compared to the MI Model group, leading to a simultaneous decrease in the protein expression levels of alpha-smooth muscle actin (α-SMA), Collagen I (Figure 3C-D).

**Figure 3.**
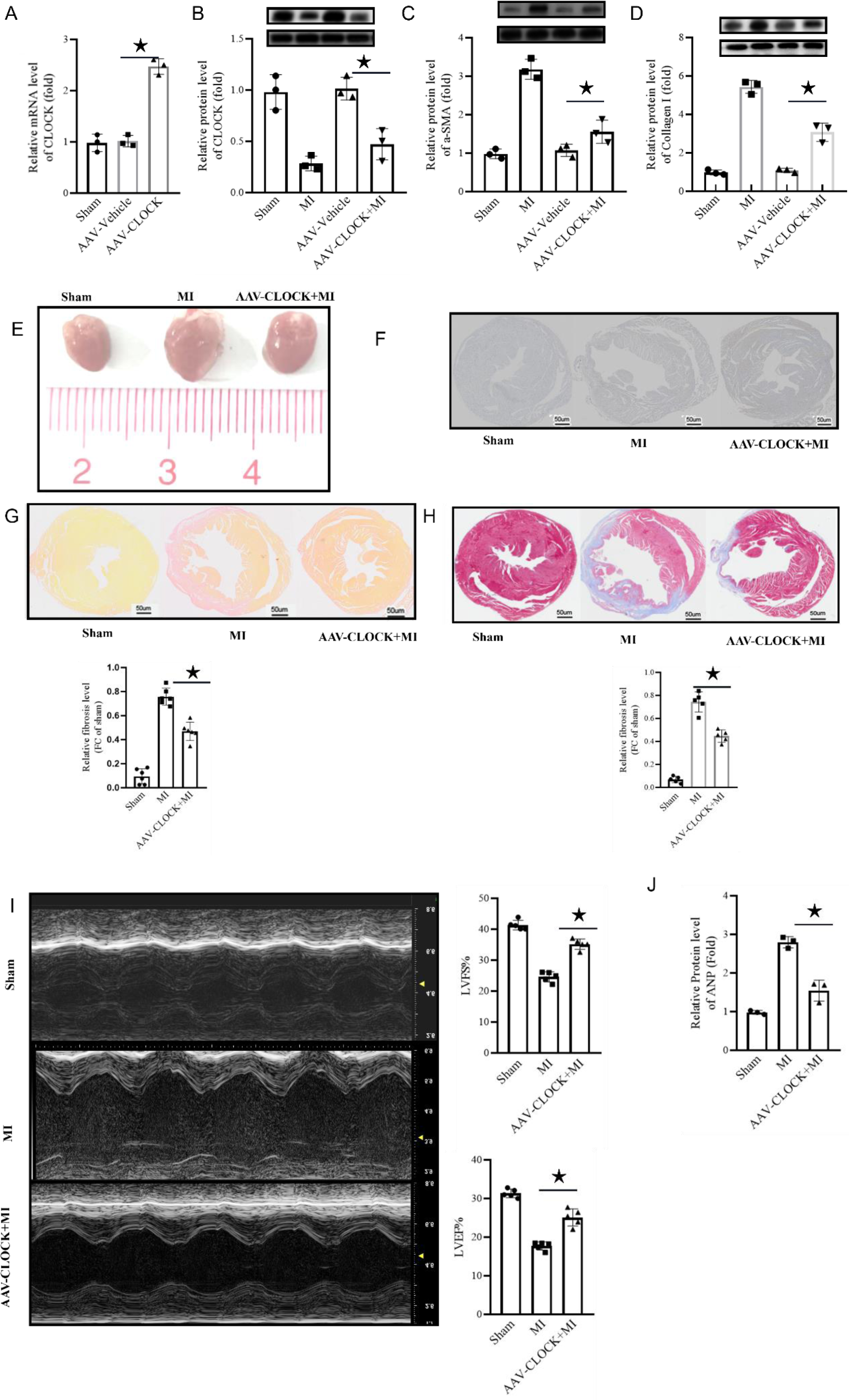
Overexpression of CLOCK in vivo reduced the severity of cardiac fibrosis and ameliorated cardiac dysfunction after myocardial infarction in mice. A: Serotype 9 adenotype-associated virus AAV carrying CLOCK overexpression plasmid, specifically overexpressed CLOCK in mouse heart, and CLOCK protein expression level were determined by western blot.N = 3; *P<0.05, ***P<0.01。 B-D: CLOCK was specifically overexpressed in mice, and a model of cardiac fibrosis was constructed. western blot detected the expression of A-SMA and Collagen I in cardiac fibrosis, N = 3; * P <0.05, * * * P <0.01. E-G: 4 weeks after the AMI or sham surgery, E: Volume comparison of the gross heart structure of the four mouse groups.F: HE staining results show the pathological comparison of mouse myocardial tissues in the four groups.G: Masson staining to assess myocardial fibrosis changes after AMI, myocardial tissue (red); collagen fiber tissue (blue), collagen volume fraction (Collagen volume fraction, CVF) to evaluate the degree of myocardial fibrosis in mice.H: Evaluation of myocardial fibrosis after AMI, collagen fiber tissue (red), collagen volume fraction (Collagen volume fraction, CVF) to evaluate the degree of myocardial fibrosis in mice.I: mice underwent cardiac ultrasound testing at week 4 after AMI or sham surgery, and cardiac function was assessed by left ventricular ejection fraction (LVEF%) and LV short axis shortening rate (LVFS%).J: qRT-PCR analysis of the mRNA levels of ANP and BNP in the hearts of the four mouse groups.

The beneficial effects of CLOCK overexpression on cardiac histopathological alterations following acute myocardial infarction (AMI) were also observed. The MI group of WT mice significantly increased the heart volume compared with the Sham group, while the CLOCK overexpression group decreased the MI group compared with the Sham group (Figure 3E). The myocardial histopathological changes after AMI were assessed by HE staining. In Sham group, there had normal myocardial structure, regular muscle fibers, clear texture and uniform myofilament clearance. In the Model group, the myocardial structure was disordered, with thick muscle filaments, waveform change and unclear texture. The myocardial histopathological changes were significantly improved in the CLOCK overexpression group (Figure 3F).

Myocardial fibrosis was assessed post-AMI using Masson staining, which revealed the presence of blue collagen fibers and red cardiac muscle fibers. The MI Model group exhibited a significant abundance of collagen fibers within the myocardial tissue, whereas CLOCK overexpression noticeably reduced collagen deposition. Collagen volume fraction (CVF) was quantified using ImageJ software, revealing a significant increase in CVF in the MI Model group compared to the Sham group (*p* <0.01). Remarkably, the CLOCK overexpression group exhibited a substantial reduction in CVF compared to the MI Model group (*p* <0.05) (Figure 3G-H).

These findings strongly suggest that CLOCK overexpression has the ability to alleviate myocardial fibrosis and mitigate the histopathological damage inflicted upon the myocardium after acute myocardial infarction. At 4 weeks post-acute myocardial infarction, mice underwent cardiac echocardiography examinations. The results of the evaluation demonstrated a substantial improvement (*p* <0.01) in LVEF and LVFS in the CLOCK overexpression model group compared to the MI Model group (Figure 3I). These observations suggest that CLOCK had the potential to significantly ameliorate the cardiac dysfunction induced by AMI. Subsequently, serum BNP levels were measured in mice, revealing a notable elevation in the MI Model group in comparison to the Sham group (*p* <0.01). Conversely, the CLOCK overexpression model group exhibited a significant reduction in serum ANP content compared to the Model group (*p* <0.01) (Figure 3J). Taken together, these findings indicate that CLOCK has the capability to effectively attenuate heart failure resulting from AMI and enhance overall cardiac function.

### CLOCK Suppresses differentiation of cardiac fibroblasts into myofibroblasts through Downregulation of Smad 3

Further investigation was conducted to explore the impact of alterations in CLOCK expression levels on the differentiation processf cardiac fibroblasts (CFs). Our research revealed the regulatory influence exerted by CLOCK on Smad 3 expression. Notably, the protein expression of Smad 3 exhibited an increase in cardiac specimens obtained from both the post-myocardial infarction (MI) model and myofibroblasts induced by AngII (Figure 4A-B). To investigate this phenomenon further, we utilized adenovirus-mediated overexpression of CLOCK or intervention with CLOCK si-RAN in cardiac fibroblasts-, Real-time PCR analysis demonstrated that Smad 3 is subject to transcriptional regulation by CLOCK in vitro using primary mouse cardiac fibroblasts. Specifically, CLOCK overexpression resulted in a decrease in Smad 3 mRNA expression- (Figure 4E). Consistent with these findings, overexpression of CLOCK resulted in a decrease in Smad 3 protein expression in cardiac fibroblasts, whereas knockdown of CLOCK led to a significant increase in Smad 3 protein levels (Figure 4C-D). These observations indicate that CLOCK represses Smad 3 expression in myofibroblasts involved in AngII-induced myocardial fibroblast differentiation, and reciprocally, Smad 3 negatively modulates CLOCK activity. To further validate the role of Smad 3 in CLOCK-mediated inhibition of cardiac fibroblast differentiation into myofibroblasts, we performed simultaneous knockdown of CLOCK and Smad 3 using si-CLOCK and si-Smad 3 in angiotensin-induced cell differentiation of cardiac fibroblasts. The knockdown of CLOCK effectively counteracted the promotion of mouse myocardial fibroblast differentiation into myofibroblasts by Smad 3. Western Blot analysis elucidated thatin the context of angiotensin-induced cell differentiation, the combined suppression of CLOCK and Smad 3 resulted in decreased Smad 3 mRNA expression and CLOCK protein expression (Figure 4F-G), leading to lower expression levels of SMA and Collagen I (Figure 4H-I). These findings provide further evidence for the enhanced differentiation of cardiac fibroblasts and confirm that CLOCK knockdown expression promotes the differentiation of fibroblast cardiomyocytes into myofibroblasts, contingent on the presence of Smad 3. Thus, our study reinforces the essential role of Smad 3 in CLOCK’s inhibition of myocardial fibroblast differentiation and unveils potential therapeutic targets for attenuating cardiac fibrosis.

**Figure 4.**
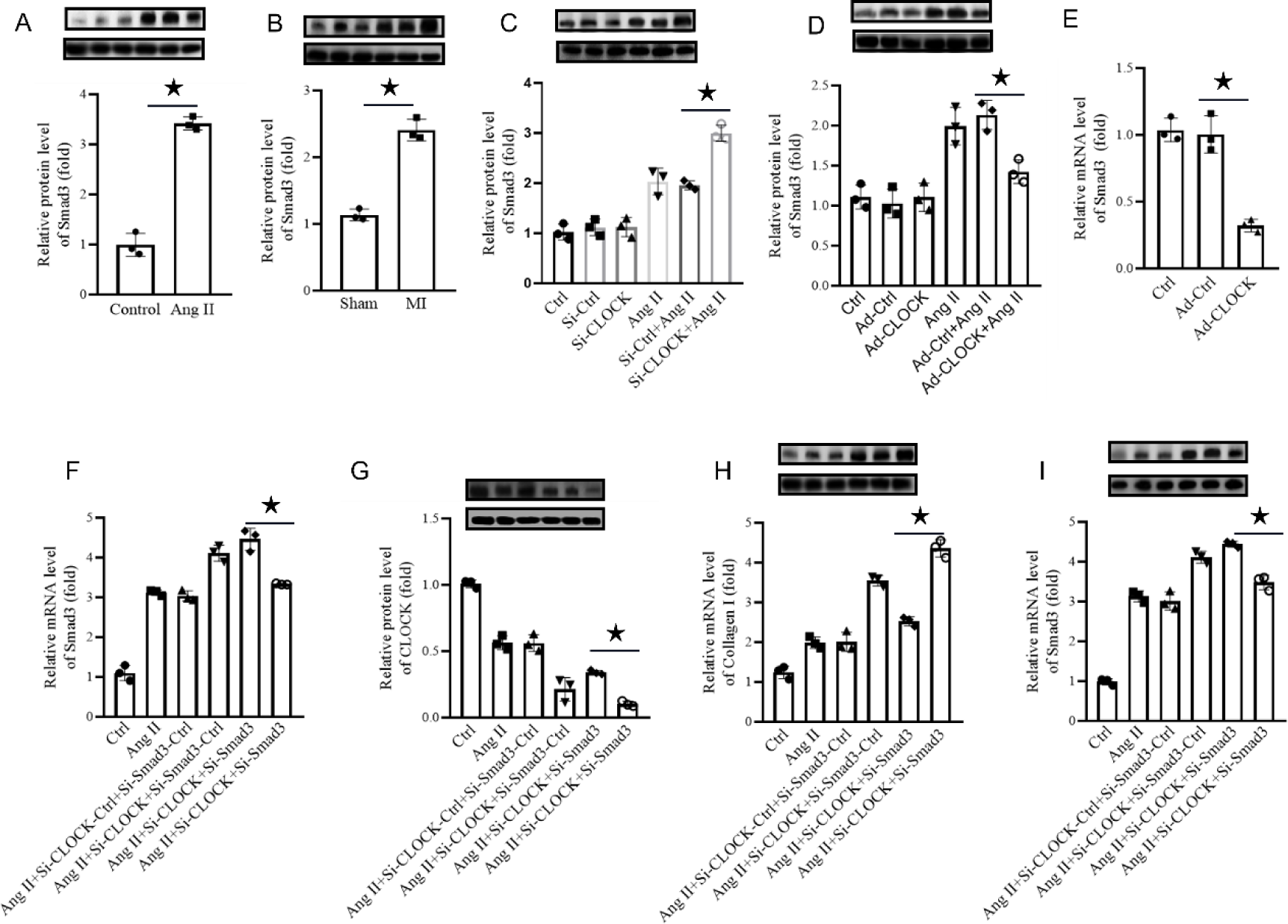
CLOCK suppresses the cellular phenotypic differentiation of inhibitory cardiac fibroblasts by regulating Smad 3 expression and function. A: In the mouse cardiac fibrosis model, the protein expression of Smad 3 was determined by western blot, N = 3; * P <0.05, * * * P <0.01. B AngII Cardiac fibroblasts were induced to differentiate into myofibroblasts, and the protein expression of Smad 3 was determined by western blot, N = 3; * P <0.05, * * * P <0.01. C CLOCK overexpression of adenovirus or CLOCK si-RAN was used to intervene in cardiac fibroblasts, and Smad3 mRNA expression was downregulated by Real-time PCR. D: Ang induced differentiation of myocardial fibroblasts, lentivirus mediated CLOCK overexpression, and A-SMA protein expression level by western blot, N = 3; * P <0.05, * * * P <0.01. E: CLOCK si-RAN intervention in cardiac fibroblasts, and a-SMA protein expression levels were measured by western blot, N = 3; * P <0.05, * * * P <0.01. F-I:Administration si-Smad 3 inhibited Smad 3 expression in cardiac fibroblasts, while downregulated CLOCK levels with si-RNA.F: mRNA expression levels of Smad 3, G-I Protein expression levels of CLOCK, Collagen I and Smad3 were detected by Western Blot; N=3; * P <0.05; * * * P <0.01.

### CLOCK binds to the Smad 3 promoter and represses transcriptional activity

In a concerted effort to elucidate the relationship between CLOCK function and Smad 3, CLOCK, identified as a transcription factor, was probed for target gene association using the Jaspar database (Figure 5A). Within the Smad 3 promoter region, seven noteworthy sequence clusters surfaced, exhibiting significant prediction scores indicative of CLOCK binding potential (Figure 5B). Employing the unique properties of transcription factors binding to DNA sequences, primers were crafted specific to these three predominant sites. Chromatin immunoprecipitation assays (ChIP-qPCR) were later conducted, leveraging a CLOCK-specific antibody. This aimed to quantify the genomic DNA association in cardiac fibroblast cells across three distinct groups: the Control, Ad-CLOCK-eGFR, and AD-CLOCK cohorts.

**Figure 5.**
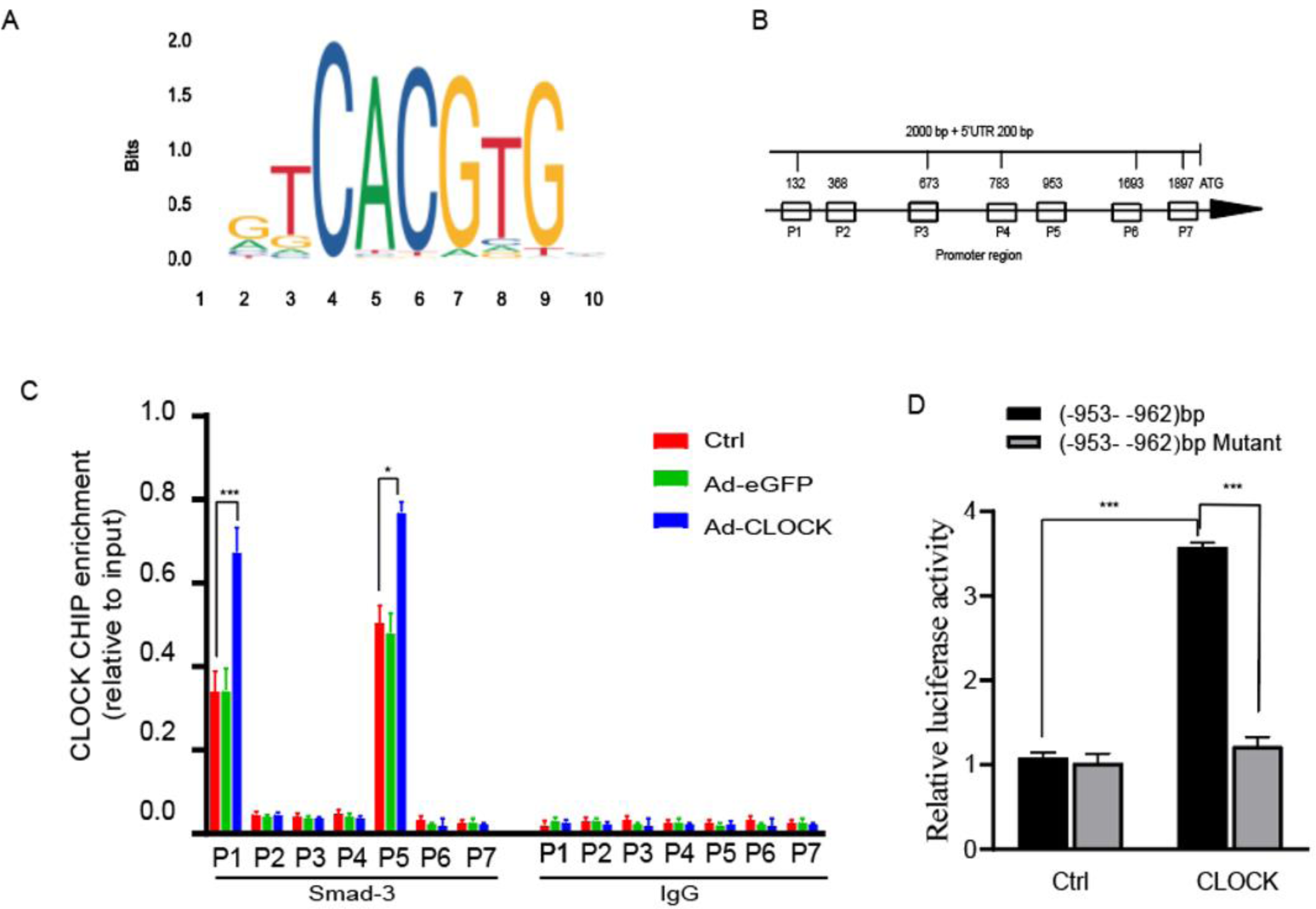
CLOCK binds to Smad3 and represses its transcriptional activity. A:Schematic diagram showing that the putative binding sites between CLOCK and Smad3 promoter by the JASPAR database.B:The binding-motif analysis showed five putative binding sites for CLOCK on the Smad3 promoter. C:Over-expression of Endogenous CLOCK by adenovirus. Chromatin immunoprecipitation (ChIP) was performed in Cardiac muscle fibroblasts of mice. Enrichment was normalized with input, using anti-CLOCK antibody and IgG ChIP as a negative control. And then showed that CLOCK bound to −953-bp Smad3 promoter region containing the two E-boxes. Five primers, P1–P7, were designed to cover the 2000 bps of HIF-1A promoter region. D:A mouse Smad3 luciferase reporter gene plasmid for the luciferase reporter gene analysis.293T Cells were transfected with the pcDNA3.1 expression plasmid and the pGL4.10-luciferase reporter plasmid. Mutation of these two E-boxes in the −953-bp Smad3 promoter disrupted the effect of binding of CLOCK and the change of the Smad3 promoter activity. All experiments were performed with at least 3 biological replicates.Data are presented as mean ± SEM; *p < 0.05, ***p < 0.001.

Intriguingly, within the Smad 3 promoter domain, the anticipated binding sites, notably the −953 bp to −962 bp intervals, manifested a marked amplification in CLOCK1 and Smad 3 DNA fragment binding. An overabundance of CLOCK1 further augmented the DNA binding presence (Figure 5C). T Such observations robustly suggest a modulatory role of CLOCK1 in Smad 3 transcription, exerted through direct gene targeting.

In an extension to the primary observations, dual-fluorophore reporter constructs were innovated for subsequent fluorophase reporter assays. The interplay between the pcDNA3.1CLOCK transcription factor expression vector and the pGL4.10-luciferaseSmad3 reporter vector was examined post-co-transfection into 293T cells. Remarkably, post a 48-hour transfection window, the fluorescence metrics were assessed, conclusively pointing to a potent inhibition of Smad 3 transcription due to CLOCK binding. Further experimentation revealed that deletion mutations of the two e-boxes residing within the −82bp and −568bp regions of the Smad 3 promoters disrupted the CLOCK binding sites, instigating an upsurge in the luciferase activity of the Smad 3 promoter (Figure 5D). Empirical evidence provided by the dual luciferase reporter assay underscored a compelling interaction between CLOCK and the Smad 3 promoter, which in turn muted the transcriptional activity of the Smad 3 gene. This interaction was found to exert a downward influence on the protein expression level of Smad 3, orchestrating the conversion of myocardial fibroblasts into myofibroblasts in a tightly regulated manner.

## Discussion

In our current exploration, the novel CLOCK-Smad3 pathway, a master regulator in the progression of myocardial fibrosis, has been elucidated. Through both in vivo and in vitro studies, we have discerned that CLOCK inhibits the transformation of cardiac fibroblasts into myofibroblasts, a critical process in the onset of cardiac fibrosis following acute myocardial infarction (AMI). Preceding scientific undertakings have emphasized the central function of Smad3 signaling within the pathological underpinnings of myocardial fibrosis. Mechanistic insights from our study reveal that CLOCK’s binding to the Smad3 promoter region functions strategically, suppressing its transcriptional activation. This mechanism results in a marked downregulation of Smad3 protein expression, impeding fibroblast-cardiomyocyte differentiation, and ultimately leading to a diminution in collagen deposition and a decrease in myofibroblast infiltration within the myocardial tissue post-MI. Thus, CLOCK is identified as an essential modulator in reducing collagen accumulation and ameliorating cardiac function in murine models subsequent to MI.

Furthermore, a complex interplay emerges between cardiac fibrosis and the malign influences of oxidative stress and elevated triglyceride concentrations. The pernicious impact of high triglyceride levels manifests in their ability to incite oxidative stress, thereby unsettling the fragile balance of redox homeostasis within cardiomyocytes. This oxidative distress, in conjunction with aberrant triglyceride metabolism, masterminds the inception and advancement of cardiac fibrosis through the induction of inflammatory responses and the engagement of signaling pathways intimately associated with the fibrotic transformation.

Extensive evidence supports the existence of rhythmic oscillation in the circadian regulation of cellular redox function, and changes in the cellular redox status significantly influence the CLOCK mechanism^51^. Importantly, oxidative stress can initiate the expression of biological rhythms, thereby modulating the activation of antioxidant transcription factors and promoting processes like autophagy and mitochondrial biogenesis^52^. The observation of circadian oscillations in H_2_O_2_ levels within mammalian cells and mouse liver confirms the direct regulation of CLOCK circadian function through cysteine oxidation^53^. Disruption of the circadian rhythm by the CLOCK gene leads to rhythmic variations in the levels of free fatty acids (FFA) and glycerol in the bloodstream of mice, resulting in the accumulation of triglycerides (TG) and subsequent enlargement adipocytes^54^. Notably, the CLOCK gene orchestrates the differential expression of regulatory genes, including those involved in adipogenesis, such as fatty acid binding protein 9 (FABP 9) and glycerol kinase 2 (GK 2)^55^,.

Lack of biological rhythm genes impairs adaptive vascular remodeling and increases collagen deposition in the medial layer of blood vessels in mice. Recent discoveries indicate a significant decrease in the oscillation expression of the biological CLOCK within the ischemic heart region. Acute ischemic events rapidly suppress circadian CLOCK genes expression in the ischemic region of the heart, potentially contributing to post-ischemic cardiac dysfunction and development of myocardial infarction^56^. In our groundbreaking study, we observed a substantial reduction in the expression of CLOCK protein in both in vivo and in vitro models of cardiac fibrosis. However, the specific association between CLOCK and cardiac fibrosis remains unclear, prompting further research to uncover the novel functionalities of CLOCK in the context of cardiac fibrosis.

In this investigation, we significantly reduced the presence of cardiac fibrosis in a murine model by overexpressing CLOCK using adeno-associated virus (AAV), compared to the model group with myocardial infarction. Additionally, CLOCK overexpression led to decreased expression levels of characteristic proteins associated with cardiac fibroblasts, such as α-SMA, type I, and type III collagen. This suggests that CLOCK also inhibits the differentiation of cardiac fibroblasts into myofibroblasts induced by Angiotensin (Ang), as observed in vitro. These findings imply that CLOCK functions as a protective regulator in the realm of cardiac fibrosis, specifically inhibiting the transdifferentiation of myocardial fibroblasts and underscoring its significance in this process. Several previous studies have elucidated the complex interplay between cardiac fibroblasts (CFs) and cardiomyocytes in the context of myocardial tissue injury. Following such injury, CFs trigger the secretion of profibrotic growth factors by cardiomyocytes^57,58^, setting in motion a cascade involving multiple profibrotic factors^59,60^. This intricate interplay ultimately leads to the upregulation of type I and type III collagen genes, as well as the promotion of fibrillary collagen and integrin gene expression, thereby driving the progression of myocardial fibrosis (MF)^47,61,62^. The release of pro-fibrotic growth factors and cytokines by CFs engages with their respective receptors, activating a multitude of signaling pathways and transcription factors, including but not limited to Smad, MAPK, Akt, and nuclear factor-kB (NF-kB). These extensively activated signaling pathways, in synergy with transcription factors, facilitate the transdifferentiation of CFs into myofibroblasts. Furthermore, the growth factors and cytokines secreted by CFs or other cellular entities can reciprocally impact both CFs and cardiomyocytes, creating a positive feedback loop that amplifies the fibrotic response.

Smad 3, a distinguished member of the renowned Smads family of proteins, plays a pivotal role in the complex realm of signaling modulated by the TGF-β superfamily^34,63^. Its indispensable contribution to the precise orchestration of cardiac fibrosis cannot be overemphasized. In the context of fibrosis, Smad 3 transcends its ordinary state and becomes hyperactivated, enabling its direct interaction with numerous promoters within the collagen matrix^64^. Remarkably, mice lacking Smad 3 demonstrated enhanced protection against the perils of fibrosis in various cardiovascular conditions^33,34,63,65^. The absence of Smad 3 acts as a formidable deterrent against collagen deposition in the scar^33^, thereby mitigating the detrimental effects of expansion remodeling and diastolic dysfunction following myocardial infarction. Additionally, targeted inhibition of Smad 3 effectively counteracts the insidious influence of Angiotensin II-induced cardiovascular hypertension^65^. It has also been convincingly demonstrated that the disruption of the Smad 3 gene directly interferes with the fibroblast responses mediated by TGF-β 1. Consequently, direct inhibition of Smad 3 emerges as an intriguing prospect, representing as a potential therapeutic target in battle against cardiac fibrosis.

Our investigation has uncovered a fascinating revelation: the attenuation of expression plasmid via Smad 3 depletion effectively undermines the protective capabilities conferred by CLOCK in vitro. This evidence raises the question of whether a reciprocal regulation between CLOCK and Smad 3 is involved in the intricate machinery governing cardiac fibrosis. The interplay between CLOCK and Smad 3 warrants further exploration, as it may shed light on novel mechanisms underlying the pathogenesis of cardiac fibrosis and potentially unveil new therapeutic avenues for its prevention and treatment. In an outstanding achievement of scientific inquiry, our study has successfully unraveled the profound influence of CLOCK in suppressing the transcriptional expression of Smad 3, leading to significant changes in the translation and synthesis of Smad 3 protein. These remarkable findings provide a solid foundation for proposing that CLOCK represents a cutting-edge therapeutic approach in the ongoing battle against cardiac fibrosis. The newfound understanding of CLOCK’s capability to modulate Smad 3 expression opens up innovative avenues for developing novel treatments and interventions aimed at combating the relentless progression of cardiac fibrosis.

---

**Supplementary Table 1.**
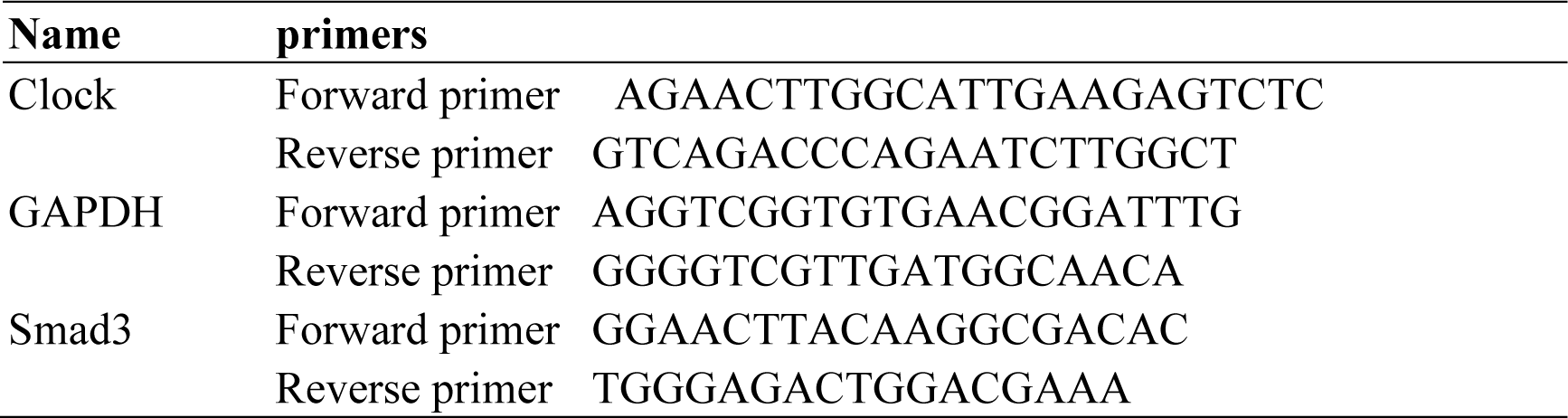
Sequences of the PCR primers.

**Supplementary Table 2.**
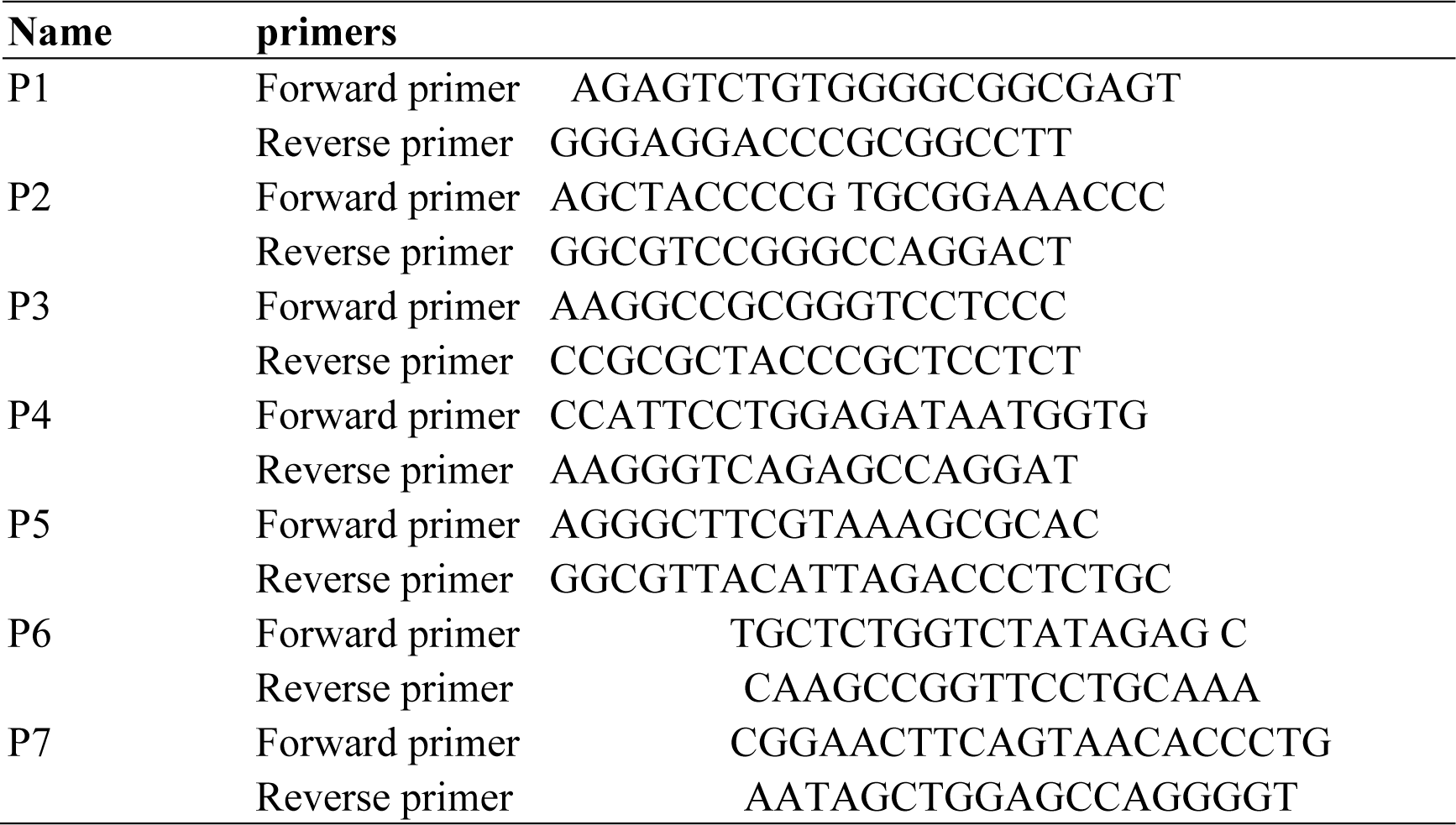
Sequences of the PCR and mutagenic primers used for ChIP-qPCR.

**Supplementary Figure 1.**
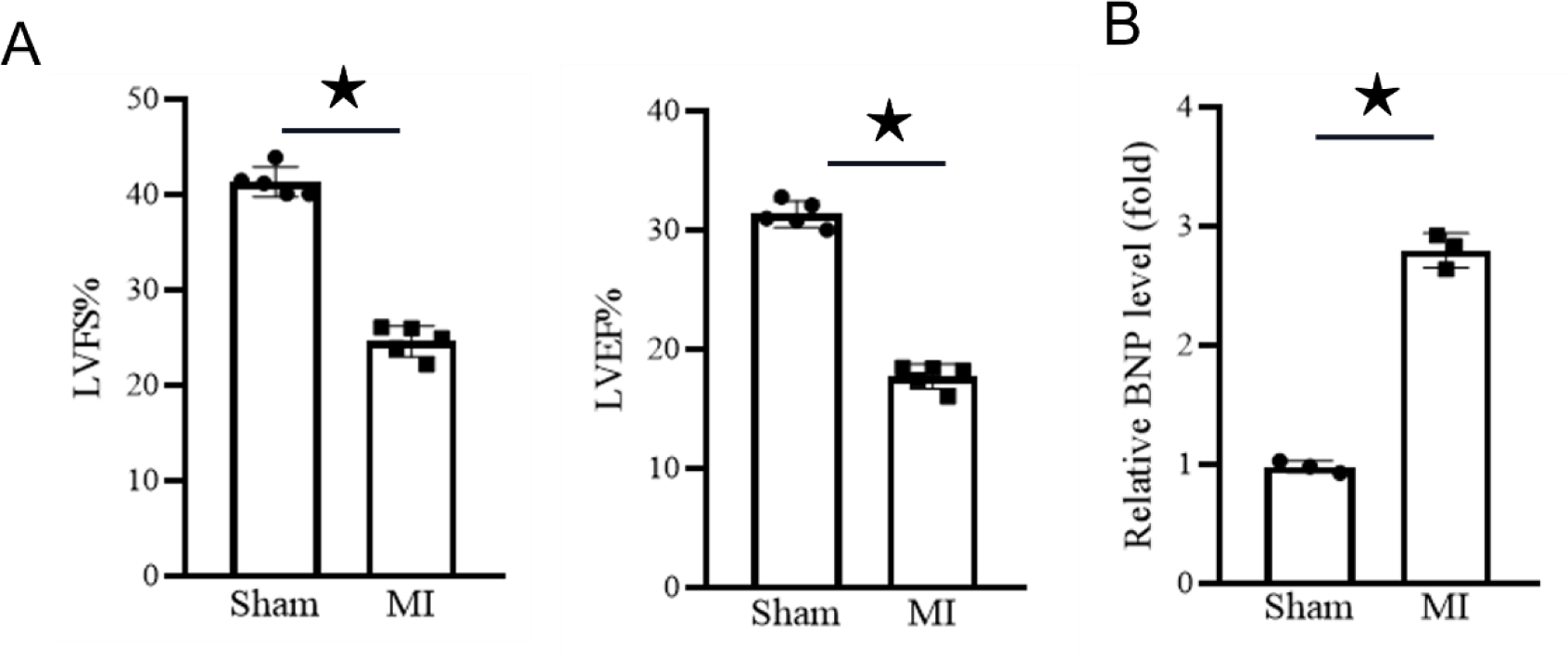
A:After serum shock for the indicated times, CLOCK mRNA levels in cardiac fibroblast cells were measured by quantitative PCR; B:Ang-induced differentiation of cardiac muscle fibroblasts into myofibroblasts, CLOCK mRNA levels were measured by quantitative PCR; N = 3, ★P<0.05.

**Supplementary Figure 2.**
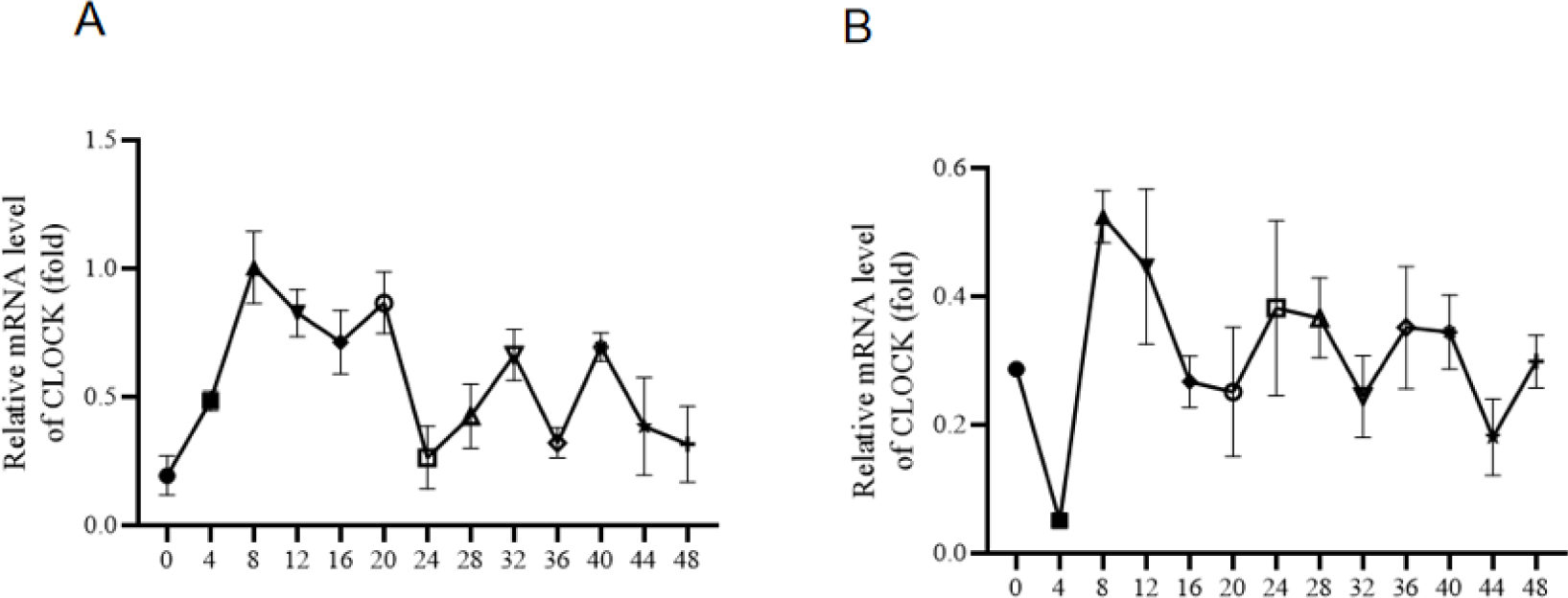
A: CLOCK circadian mRNA expression of control group. After serum shock for the indicated times, mRNA levels in cardiac muscle fibroblasts were measured by quantitative PCR. B: After serum shock for the indicated times (hour), 10^−6^M AngII stimulated mouse primary cardiac myocardial fibroblasts induced differentiation into myofibroblasts, mRNA levels of CLOCK were measured by quantitative PCR.

## References

1. Levitan BM, Manning JR, Withers CN, Smith JD, Shaw RM, Andres DA, Sorrell VL, Satin J. Rad-deletion Phenocopies Tonic Sympathetic Stimulation of the Heart. J Cardiovasc Transl Res. 2016;9:432–444. doi: 10.1007/s12265-016-9716-y

2. Yin L, Liu MX, Li W, Wang FY, Tang YH, Huang CX. Over-Expression of Inhibitor of Differentiation 2 Attenuates Post-Infarct Cardiac Fibrosis Through Inhibition of TGF-beta1/Smad3/HIF-1alpha/IL-11 Signaling Pathway. Front Pharmacol. 2019;10:1349. doi: 10.3389/fphar.2019.01349

3. Chen X, Li H, Wang K, Liang X, Wang W, Hu X, Huang Z, Wang Y. Aerobic Exercise Ameliorates Myocardial Inflammation, Fibrosis and Apoptosis in High-Fat-Diet Rats by Inhibiting P2X7 Purinergic Receptors. Front Physiol. 2019;10:1286. doi: 10.3389/fphys.2019.01286

4. Frangogiannis NG. The inflammatory response in myocardial injury, repair, and remodelling. Nat Rev Cardiol. 2014;11:255–265. doi: 10.1038/nrcardio.2014.28

5. Karpov AA, Udalova DV, Pliss MG, Galagudza MM. Can the outcomes of mesenchymal stem cell-based therapy for myocardial infarction be improved? Providing weapons and armour to cells. Cell Prolif. 2017;50. doi: 10.1111/cpr.12316

6. Schirone L, Forte M, Palmerio S, Yee D, Nocella C, Angelini F, Pagano F, Schiavon S, Bordin A, Carrizzo A, et al. A Review of the Molecular Mechanisms Underlying the Development and Progression of Cardiac Remodeling. Oxid Med Cell Longev. 2017;2017:3920195. doi: 10.1155/2017/3920195

7. Querejeta R, Lopez B, Gonzalez A, Sanchez E, Larman M, Martinez Ubago JL, Diez J. Increased collagen type I synthesis in patients with heart failure of hypertensive origin: relation to myocardial fibrosis. Circulation. 2004;110:1263–1268. doi: 10.1161/01.CIR.0000140973.60992.9A

8. Shinde AV, Frangogiannis NG. Mechanisms of Fibroblast Activation in the Remodeling Myocardium. Curr Pathobiol Rep. 2017;5:145–152. doi: 10.1007/s40139-017-0132-z

9. Grigorian Shamagian L, Madonna R, Taylor D, Climent AM, Prosper F, Bras-Rosario L, Bayes-Genis A, Ferdinandy P, Fernandez-Aviles F, Izpisua Belmonte JC, et al. Perspectives on Directions and Priorities for Future Preclinical Studies in Regenerative Medicine. Circ Res. 2019;124:938–951. doi: 10.1161/CIRCRESAHA.118.313795

10. Tanaka R, Umemura M, Narikawa M, Hikichi M, Osaw K, Fujita T, Yokoyama U, Ishigami T, Tamura K, Ishikawa Y. Reactive fibrosis precedes doxorubicin-induced heart failure through sterile inflammation. ESC Heart Fail. 2020;7:588–603. doi: 10.1002/ehf2.12616

11. Biswas S, Thomas AA, Chakrabarti S. LncRNAs: Proverbial Genomic “Junk” or Key Epigenetic Regulators During Cardiac Fibrosis in Diabetes? Front Cardiovasc Med. 2018;5:28. doi: 10.3389/fcvm.2018.00028

12. Videira RF, da Costa Martins PA. Non-coding RNAs in Cardiac Intercellular Communication. Front Physiol. 2020;11:738. doi: 10.3389/fphys.2020.00738

13. Guo X, Yan F, Li J, Zhang C, Bu P. SIRT3 attenuates AngII-induced cardiac fibrosis by inhibiting myofibroblasts transdifferentiation via STAT3-NFATc2 pathway. Am J Transl Res. 2017;9:3258–3269.

14. Jeong D, Lee MA, Li Y, Yang DK, Kho C, Oh JG, Hong G, Lee A, Song MH, LaRocca TJ, et al. Matricellular Protein CCN5 Reverses Established Cardiac Fibrosis. J Am Coll Cardiol. 2016;67:1556–1568. doi: 10.1016/j.jacc.2016.01.030

15. Jugdutt BI. Ventricular remodeling after infarction and the extracellular collagen matrix: when is enough enough? Circulation. 2003;108:1395–1403. doi: 10.1161/01.CIR.0000085658.98621.49

16. Wang Y, Ouyang M, Wang Q, Jian Z. MicroRNA-142-3p inhibits hypoxia/reoxygenation-induced apoptosis and fibrosis of cardiomyocytes by targeting high mobility group box 1. Int J Mol Med. 2016;38:1377–1386. doi: 10.3892/ijmm.2016.2756

17. Zhou T, Wang Z, Fan J, Chen S, Tan Z, Yang H, Yin Y. Angiotensin-converting enzyme-2 overexpression improves atrial remodeling and function in a canine model of atrial fibrillation. J Am Heart Assoc. 2015;4:e001530. doi: 10.1161/JAHA.114.001530

18. Xin JJ, Dai QF, Lu FY, Zhao YX, Liu Q, Cui JJ, Xu DS, Bai WZ, Jing XH, Gao JH, Yu XC. Antihypertensive and Antifibrosis Effects of Acupuncture at PC6 Acupoints in Spontaneously Hypertensive Rats and the Underlying Mechanisms. Front Physiol. 2020;11:734. doi: 10.3389/fphys.2020.00734

19. Boarescu PM, Chirila I, Bulboaca AE, Bocsan IC, Pop RM, Gheban D, Bolboaca SD. Effects of Curcumin Nanoparticles in Isoproterenol-Induced Myocardial Infarction. Oxid Med Cell Longev. 2019;2019:7847142. doi: 10.1155/2019/7847142

20. Xiao J, Sheng X, Zhang X, Guo M, Ji X. Curcumin protects against myocardial infarction-induced cardiac fibrosis via SIRT1 activation in vivo and in vitro. Drug Des Devel Ther. 2016;10:1267–1277. doi: 10.2147/DDDT.S104925

21. Sun L, Jin H, Sun L, Chen S, Huang Y, Liu J, Li Z, Zhao M, Sun Y, Tang C, et al. Hydrogen sulfide alleviates myocardial collagen remodeling in association with inhibition of TGF-beta/Smad signaling pathway in spontaneously hypertensive rats. Mol Med. 2015;20:503–515. doi: 10.2119/molmed.2013.00096

22. Atale N, Yadav D, Rani V, Jin JO. Pathophysiology, Clinical Characteristics of Diabetic Cardiomyopathy: Therapeutic Potential of Natural Polyphenols. Front Nutr. 2020;7:564352. doi: 10.3389/fnut.2020.564352

23. Uitterdijk A, Springeling T, Hermans KCM, Merkus D, de Beer VJ, Gorsse-Bakker C, Mokelke E, Daskalopoulos EP, Wielopolski PA, Cleutjens JPM, et al. Intermittent pacing therapy favorably modulates infarct remodeling. Basic Res Cardiol. 2017;112:28. doi: 10.1007/s00395-017-0616-3

24. Heldin CH, Miyazono K, ten Dijke P. TGF-beta signalling from cell membrane to nucleus through SMAD proteins. Nature. 1997;390:465–471. doi: 10.1038/37284

25. Shi Y, Massague J. Mechanisms of TGF-beta signaling from cell membrane to the nucleus. Cell. 2003;113:685–700. doi: 10.1016/s0092-8674(03)00432-x

26. Abdel-Latif A, Zuba-Surma EK, Case J, Tiwari S, Hunt G, Ranjan S, Vincent RJ, Srour EF, Bolli R, Dawn B. TGF-beta1 enhances cardiomyogenic differentiation of skeletal muscle-derived adult primitive cells. Basic Res Cardiol. 2008;103:514–524. doi: 10.1007/s00395-008-0729-9

27. Deres L, Eros K, Horvath O, Bencze N, Cseko C, Farkas S, Habon T, Toth K, Halmosi R. The Effects of Bradykinin B1 Receptor Antagonism on the Myocardial and Vascular Consequences of Hypertension in SHR Rats. Front Physiol. 2019;10:624. doi: 10.3389/fphys.2019.00624

28. Lei R, Li J, Liu F, Li W, Zhang S, Wang Y, Chu X, Xu J. HIF-1alpha promotes the keloid development through the activation of TGF-beta/Smad and TLR4/MyD88/NF-kappaB pathways. Cell Cycle. 2019;18:3239–3250. doi: 10.1080/15384101.2019.1670508

29. Liao HH, Zhu JX, Feng H, Ni J, Zhang N, Chen S, Liu HJ, Yang Z, Deng W, Tang QZ. Myricetin Possesses Potential Protective Effects on Diabetic Cardiomyopathy through Inhibiting IkappaBalpha/NFkappaB and Enhancing Nrf2/HO-1. Oxid Med Cell Longev. 2017;2017:8370593. doi: 10.1155/2017/8370593

30. Zhou J, Zhong DW, Wang QW, Miao XY, Xu XD. Paclitaxel ameliorates fibrosis in hepatic stellate cells via inhibition of TGF-beta/Smad activity. World J Gastroenterol. 2010;16:3330–3334. doi: 10.3748/wjg.v16.i26.3330

31. Shang L, Zhang L, Shao M, Feng M, Shi J, Dong Z, Guo Q, Xiaokereti J, Xiang R, Sun H, et al. Elevated beta1-Adrenergic Receptor Autoantibody Levels Increase Atrial Fibrillation Susceptibility by Promoting Atrial Fibrosis. Front Physiol. 2020;11:76. doi: 10.3389/fphys.2020.00076

32. Jiang C, Xie N, Sun T, Ma W, Zhang B, Li W. Xanthohumol Inhibits TGF-beta1-Induced Cardiac Fibroblasts Activation via Mediating PTEN/Akt/mTOR Signaling Pathway. Drug Des Devel Ther. 2020;14:5431–5439. doi: 10.2147/DDDT.S282206

33. Bujak M, Ren G, Kweon HJ, Dobaczewski M, Reddy A, Taffet G, Wang XF, Frangogiannis NG. Essential role of Smad3 in infarct healing and in the pathogenesis of cardiac remodeling. Circulation. 2007;116:2127–2138. doi: 10.1161/CIRCULATIONAHA.107.704197

34. Dobaczewski M, Bujak M, Li N, Gonzalez-Quesada C, Mendoza LH, Wang XF, Frangogiannis NG. Smad3 signaling critically regulates fibroblast phenotype and function in healing myocardial infarction. Circ Res. 2010;107:418–428. doi: 10.1161/CIRCRESAHA.109.216101

35. Kong P, Shinde AV, Su Y, Russo I, Chen B, Saxena A, Conway SJ, Graff JM, Frangogiannis NG. Opposing Actions of Fibroblast and Cardiomyocyte Smad3 Signaling in the Infarcted Myocardium. Circulation. 2018;137:707–724. doi: 10.1161/CIRCULATIONAHA.117.029622

36. Chen B, Huang S, Su Y, Wu YJ, Hanna A, Brickshawana A, Graff J, Frangogiannis NG. Macrophage Smad3 Protects the Infarcted Heart, Stimulating Phagocytosis and Regulating Inflammation. Circ Res. 2019;125:55–70. doi: 10.1161/CIRCRESAHA.119.315069

37. Ma ZG, Yuan YP, Zhang X, Xu SC, Wang SS, Tang QZ. Piperine Attenuates Pathological Cardiac Fibrosis Via PPAR-gamma/AKT Pathways. EBioMedicine. 2017;18:179–187. doi: 10.1016/j.ebiom.2017.03.021

38. Huang S, Chen B, Su Y, Alex L, Humeres C, Shinde AV, Conway SJ, Frangogiannis NG. Distinct roles of myofibroblast-specific Smad2 and Smad3 signaling in repair and remodeling of the infarcted heart. J Mol Cell Cardiol. 2019;132:84–97. doi: 10.1016/j.yjmcc.2019.05.006

39. King DP, Zhao Y, Sangoram AM, Wilsbacher LD, Tanaka M, Antoch MP, Steeves TD, Vitaterna MH, Kornhauser JM, Lowrey PL, et al. Positional cloning of the mouse circadian clock gene. Cell. 1997;89:641–653. doi: 10.1016/s0092-8674(00)80245-7

40. Gekakis N, Staknis D, Nguyen HB, Davis FC, Wilsbacher LD, King DP, Takahashi JS, Weitz CJ. Role of the CLOCK protein in the mammalian circadian mechanism. Science. 1998;280:1564–1569. doi: 10.1126/science.280.5369.1564

41. Antoch MP, Song EJ, Chang AM, Vitaterna MH, Zhao Y, Wilsbacher LD, Sangoram AM, King DP, Pinto LH, Takahashi JS. Functional identification of the mouse circadian Clock gene by transgenic BAC rescue. Cell. 1997;89:655–667. doi: 10.1016/s0092-8674(00)80246-9

42. Bray MS, Shaw CA, Moore MW, Garcia RA, Zanquetta MM, Durgan DJ, Jeong WJ, Tsai JY, Bugger H, Zhang D, et al. Disruption of the circadian clock within the cardiomyocyte influences myocardial contractile function, metabolism, and gene expression. Am J Physiol Heart Circ Physiol. 2008;294:H1036–1047. doi: 10.1152/ajpheart.01291.2007

43. Alibhai FJ, Tsimakouridze EV, Reitz CJ, Pyle WG, Martino TA. Consequences of Circadian and Sleep Disturbances for the Cardiovascular System. Can J Cardiol. 2015;31:860–872. doi: 10.1016/j.cjca.2015.01.015

44. Podobed P, Pyle WG, Ackloo S, Alibhai FJ, Tsimakouridze EV, Ratcliffe WF, Mackay A, Simpson J, Wright DC, Kirby GM, et al. The day/night proteome in the murine heart. Am J Physiol Regul Integr Comp Physiol. 2014;307:R121–137. doi: 10.1152/ajpregu.00011.2014

45. Podobed PS, Alibhai FJ, Chow CW, Martino TA. Circadian regulation of myocardial sarcomeric Titin-cap (Tcap, telethonin): identification of cardiac clock-controlled genes using open access bioinformatics data. PLoS One. 2014;9:e104907. doi: 10.1371/journal.pone.0104907

46. Sato H, Watanabe A, Tanaka T, Koitabashi N, Arai M, Kurabayashi M, Yokoyama T. Regulation of the human tumor necrosis factor-alpha promoter by angiotensin II and lipopolysaccharide in cardiac fibroblasts: different cis-acting promoter sequences and transcriptional factors. J Mol Cell Cardiol. 2003;35:1197–1205. doi: 10.1016/s0022-2828(03)00210-4

47. Schorb W, Booz GW, Dostal DE, Conrad KM, Chang KC, Baker KM. Angiotensin II is mitogenic in neonatal rat cardiac fibroblasts. Circ Res. 1993;72:1245–1254. doi: 10.1161/01.res.72.6.1245

48. Gao X, He X, Luo B, Peng L, Lin J, Zuo Z. Angiotensin II increases collagen I expression via transforming growth factor-beta1 and extracellular signal-regulated kinase in cardiac fibroblasts. Eur J Pharmacol. 2009;606:115–120. doi: 10.1016/j.ejphar.2008.12.049

49. Zhang R, Zhang YY, Huang XR, Wu Y, Chung AC, Wu EX, Szalai AJ, Wong BC, Lau CP, Lan HY. C-reactive protein promotes cardiac fibrosis and inflammation in angiotensin II-induced hypertensive cardiac disease. Hypertension. 2010;55:953–960. doi: 10.1161/HYPERTENSIONAHA.109.140608

50. Schnee JM, Hsueh WA. Angiotensin II, adhesion, and cardiac fibrosis. Cardiovasc Res. 2000;46:264–268. doi: 10.1016/s0008-6363(00)00044-4

51. Lananna BV, Musiek ES. The wrinkling of time: Aging, inflammation, oxidative stress, and the circadian clock in neurodegeneration. Neurobiol Dis. 2020;139:104832. doi: 10.1016/j.nbd.2020.104832

52. Sengupta S, Yang G, O’Donnell JC, Hinson MD, McCormack SE, Falk MJ, La P, Robinson MB, Williams ML, Yohannes MT, et al. The circadian gene Rev-erbalpha improves cellular bioenergetics and provides preconditioning for protection against oxidative stress. Free Radic Biol Med. 2016;93:177–189. doi: 10.1016/j.freeradbiomed.2016.02.004

53. Pei JF, Li XK, Li WQ, Gao Q, Zhang Y, Wang XM, Fu JQ, Cui SS, Qu JH, Zhao X, et al. Diurnal oscillations of endogenous H(2)O(2) sustained by p66(Shc) regulate circadian clocks. Nat Cell Biol. 2019;21:1553–1564. doi: 10.1038/s41556-019-0420-4

54. Shostak A, Meyer-Kovac J, Oster H. Circadian regulation of lipid mobilization in white adipose tissues. Diabetes. 2013;62:2195–2203. doi: 10.2337/db12-1449

55. Bray MS, Young ME. The role of cell-specific circadian clocks in metabolism and disease. Obes Rev. 2009;10 Suppl 2:6–13. doi: 10.1111/j.1467-789X.2009.00684.x

56. Kung TA, Egbejimi O, Cui J, Ha NP, Durgan DJ, Essop MF, Bray MS, Shaw CA, Hardin PE, Stanley WC, Young ME. Rapid attenuation of circadian clock gene oscillations in the rat heart following ischemia-reperfusion. J Mol Cell Cardiol. 2007;43:744–753. doi: 10.1016/j.yjmcc.2007.08.018

57. Kamo T, Akazawa H, Komuro I. Cardiac nonmyocytes in the hub of cardiac hypertrophy. Circ Res. 2015;117:89–98. doi: 10.1161/CIRCRESAHA.117.305349

58. Sadoshima J, Xu Y, Slayter HS, Izumo S. Autocrine release of angiotensin II mediates stretch-induced hypertrophy of cardiac myocytes in vitro. Cell. 1993;75:977–984. doi: 10.1016/0092-8674(93)90541-w

59. Frieler RA, Mortensen RM. Immune cell and other noncardiomyocyte regulation of cardiac hypertrophy and remodeling. Circulation. 2015;131:1019–1030. doi: 10.1161/CIRCULATIONAHA.114.008788

60. Kong P, Christia P, Frangogiannis NG. The pathogenesis of cardiac fibrosis. Cell Mol Life Sci. 2014;71:549–574. doi: 10.1007/s00018-013-1349-6

61. Crabos M, Roth M, Hahn AW, Erne P. Characterization of angiotensin II receptors in cultured adult rat cardiac fibroblasts. Coupling to signaling systems and gene expression. J Clin Invest. 1994;93:2372–2378. doi: 10.1172/JCI117243

62. Sadoshima J, Izumo S. Molecular characterization of angiotensin II--induced hypertrophy of cardiac myocytes and hyperplasia of cardiac fibroblasts. Critical role of the AT1 receptor subtype. Circ Res. 1993;73:413–423. doi: 10.1161/01.res.73.3.413

63. Huang XR, Chung AC, Yang F, Yue W, Deng C, Lau CP, Tse HF, Lan HY. Smad3 mediates cardiac inflammation and fibrosis in angiotensin II-induced hypertensive cardiac remodeling. Hypertension. 2010;55:1165–1171. doi: 10.1161/HYPERTENSIONAHA.109.147611

64. Chen SJ, Yuan W, Mori Y, Levenson A, Trojanowska M, Varga J. Stimulation of type I collagen transcription in human skin fibroblasts by TGF-beta: involvement of Smad 3. J Invest Dermatol. 1999;112:49–57. doi: 10.1046/j.1523-1747.1999.00477.x

65. Liu Z, Huang XR, Lan HY. Smad3 mediates ANG II-induced hypertensive kidney disease in mice. Am J Physiol Renal Physiol. 2012;302:F986–997. doi: 10.1152/ajprenal.00595.2011

